# Spontaneous recovery and the multiple timescales of human motor memory

**DOI:** 10.1101/2020.03.24.006023

**Authors:** Simon P. Orozco, Scott T. Albert, Reza Shadmehr

## Abstract

In numerous paradigms, from fear conditioning to motor adaptation, memory exhibits a remarkable property: acquisition of a novel behavior followed by its extinction results in spontaneous recovery of the original behavior. A current model suggests that spontaneous recovery occurs because learning is supported by two different adaptive processes: one fast (high error sensitivity, low retention), and the other slow (low error sensitivity, high retention). Here, we searched for signatures of these hypothesized processes in the commands that guided single movements. We examined human saccadic eye movements and observed that following experience of a visual error, there was an adaptive change in the motor commands of the subsequent saccade, partially correcting for the error. However, the error correcting commands were expressed only during the deceleration period. If the errors persisted, the acceleration period commands also changed. Adaptation of acceleration period commands exhibited poor sensitivity to error, but the learning was resistant to forgetting. In contrast, the deceleration period commands adapted with high sensitivity to error, and the learning suffered from poor retention. Thus, within a single saccade, a fast-like process influenced the deceleration period commands, whereas a slow-like process influenced the acceleration period commands. Following extinction training, with passage of time motor memory exhibited spontaneous recovery, as evidenced by return of saccade endpoints toward their initial adapted state. The temporal dynamics of spontaneous recovery suggested that a single saccade is controlled by two different adaptive controllers, one active during acceleration, and the other during deceleration.

**Significance statement:** A feature of memory in many paradigms is the phenomenon of spontaneous recovery: learning followed by extinction inevitably leads to reversion toward the originally learned behavior. A theoretical model posits that spontaneous recovery is a feature of memory systems that learn with two independent learning processes, one fast, and the other slow. However, there have been no direct measures of these putative processes. Here, we found potential signatures of the two independent adaptive processes during control of a single saccade. The results suggest that distinct adaptive controllers contribute to the acceleration and deceleration phases of a saccade, and that each controller is supported by a fast and a slow learning process.

## Introduction

A general property of memory is the phenomenon of spontaneous recovery. For example, in the field of classical conditioning, bees can learn to associate an odor with nectar, extending their proboscis upon presentation of the odor (Stollhoff et al., 2005). They will extinguish this response if the odor is presented without the nectar. However, following passage of time, the bees once again extend their proboscis when they are presented with the odor. In the field of fear conditioning, a stimulus can be associated with a shock, inducing fear. This fear can be extinguished if the stimulus is presented without the shock (Schiller et al., 2010). However, fear of the stimulus returns following passage of time. In the field of motor learning, people and other animals respond to a perturbation by modifying their motor commands through learning from error. If the perturbation changes direction, reversing the error vector, behavior returns to baseline. However, with passage of time the behavior spontaneously reverts back: subjects reproduce the motor commands that they had originally learned (Criscimagna-Hemminger & Shadmehr, 2008; Kojima et al., 2004; Sarwary et al., 2018; Smith et al., 2006). Similar patterns of behavior have been noted in motion adaptation (Mei et al., 2018), prism adaptation (Inoue et al., 2014), facial adaptation (Mesik et al., 2013), and contrast adaptation (Bao & Engel, 2012). The central theme in all of these learning episodes is spontaneous recovery. Why does memory exhibit spontaneous recovery?

A mathematical model (Kording et al., 2007; Smith et al., 2006) suggests that during learning, changes in behavior are supported by two adaptive processes: a fast adaptive process that has high sensitivity to error along with poor retention, and a slow adaptive process that has poor sensitivity to error along with robust retention. In this model, learning followed by extinction produces competition between the fast and the slow processes. With passage of time, the fast process decays, inducing spontaneous recovery of the previously learned behavior.

Despite the simplicity of this model, it has been difficult to test the link between spontaneous recovery and the putative fast and slow adaptive processes. In the field of motor learning, correlates of the fast and slow processes have been found in the synaptic plasticity mechanisms of single neurons or groups of neurons in the cerebellum (Casellato et al., 2015, 2015; Hall et al., 2018; Herzfeld et al., 2018; Mandelblat-Cerf et al., 2011; Yamazaki et al., 2015; Yang & Lisberger, 2014). Correlates of the fast and slow processes have also been noted in distinct regions of the cerebral cortex and the cerebellum (Cassady et al., 2018; Della-Maggiore et al., 2017; Galea et al., 2011; Hadipour-Niktarash et al., 2007; Hamel et al., 2017; Herzfeld et al., 2014; Kim et al., 2015; Landi et al., 2011; Li et al., 2011; Medina et al., 2001; Ruitenberg et al., 2018; Trewartha et al., 2014; Villalta et al., 2015; Werner et al., 2014). Finally, different memory systems (explicit and implicit) have been proposed to contribute to the two putative processes (Holland et al., 2018; McDougle et al., 2015).

In order to understand the mechanisms of spontaneous recovery, we thought it crucial to search for behavioral signatures of the putative fast and slow processes. Here, we focused on saccade adaptation and made a surprising discovery: following experience of a single error, which in theory predominantly engaged the fast process, there was an adaptive change in the motor commands of the subsequent saccade, but that change corrected for the error only during the deceleration phase of the movement. If the errors persisted for many trials, both the acceleration and deceleration period commands changed. However, whereas the acceleration period commands adapted with poor sensitivity to error, the adaptation during this period of the movement exhibited robust retention. On the other hand, whereas the deceleration period commands adapted with high sensitivity to error, the adaptation during this period exhibited poor retention. Thus, the deceleration and acceleration period commands of a single saccade differentially expressed properties of the putative fast and slow processes.

With this behavioral proxy in hand, we examined spontaneous recovery. Following learning and extinction, saccade endpoint exhibited spontaneous recovery, but importantly, saccade trajectory had trial-by-trial dynamics that were markedly different than those predicted by the current models. Rather, the results suggest a new model of motor memory in which the commands that guide a single saccade are supported by two distinct adaptive controllers, one that specializes in contributing to acceleration, and the other that specializes in contributing to deceleration. Spontaneous recovery appears to occur because each controller is supported by multiple timescales of memory.

## Materials and Method

Our goal was to ask whether there were measurable behavioral correlates of the fast and slow adaptative processes within a single movement. In order to do so, we focused on a cross-axis saccade adaptation (Chen-Harris et al., 2008; Xu-Wilson et al., 2009): in all experiments, the primary target was placed at 15°, and upon saccade initiation (on perturbed trials), the target was erased and replaced with a new target at 5° perpendicular to the direction of the original target (Fig. 1A). Thus, the primary movement direction and direction of error were perpendicular. This approach has its roots in force field reaching experiments (Brashers-Krug et al., 1996), and has the advantage that it disambiguates the primary movement commands from the commands that change because of error experience.

**Figure 1.**
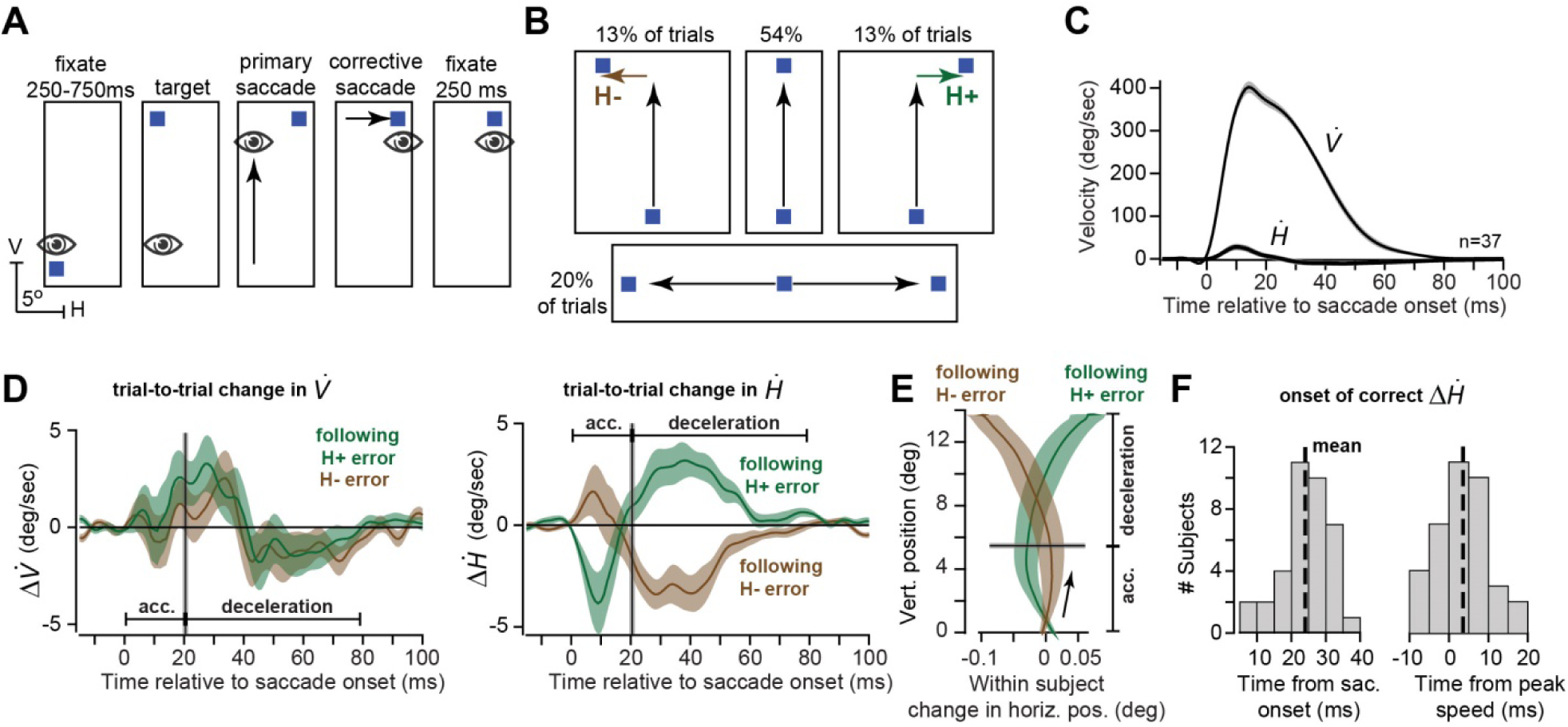
Following experience of a single error, correction on the subsequent saccade is primarily during the deceleration period. **A**. Subjects made vertical saccades and occasionally experienced a horizontal error. **B**. The structure of the various trials. **C**. Average horizontal and vertical velocity of the saccade for the target along the vertical axis. **D**. Trial-to-trial change in vertical and horizontal velocity following experience of a positive or negative horizontal endpoint error. Vertical line indicates time of peak speed, thus separating the acceleration and deceleration periods. **E**. Trial-to-trial change in saccade trajectory following experience of a positive or negative horizontal endpoint error. The endpoint correction is primarily due to commands in the deceleration period. **F**. Onset time of correction in the horizontal velocity following experience of a horizontal endpoint error. Mean onset time is about 3 ms following start of the deceleration period. Error bars are between subject SEM.

All subjects for all experiments gave written informed consent in accordance with policies approved by the Johns Hopkins School of Medicine Institutional Review Board.

### General protocol

Participants sat in a chair with their head restrained by a bite-bar made from dental putty (Coltene-Whaledent Inc. Cuyahoga Falls, Ohio, USA). They viewed a computer monitor (AGON 27” 2560×1440, 144Hz) which displayed small square targets (0.25° with 0.1° dark border). They made saccades in response to presentation of the target, and eye movements were recorded using Eyelink 1000 (S-R Research Ottawa, Ontario, CA).

The basic trial structure is shown in Fig. 1A. Each trial began with a fixation period of variable delay (randomly sampled from a uniform distribution 250-750ms), during which subjects had to maintain fixation of the target in the center of the screen within a tolerance window of ±2.5°. If they blinked or looked away early, the timer was reset and the trial repeated.

All experiments started with a baseline period during which there were no target jumps. During the baseline block, we measured saccade kinematics to targets that were randomly presented on the vertical or horizontal axes, as well as targets along various oblique directions. Vertical saccades were made to a target at (0,+15°). Horizontal saccades (20% of the trials) were made to targets at (±15,0°). Oblique saccades were made to targets at (±1,15°), (±2,15°), (±3,15°), (±4,15°), or (±5,15°).

### Experiment 1

Experimental setup is shown in Fig. 1B. We collected data from 38 subjects, but one was excluded due to poor calibration that made over 25% of trials unusable (final n=37, 26.08 ± 8.9 years old, 22 females). Rightward and leftward perturbations were randomly interleaved with control (no jump) trials so that every perturbation trial was preceded and followed by at least one control trial. We divided each saccade into an acceleration phase and a deceleration phase based on peak speed (magnitude of the velocity vector), which was largely dominated by the vertical component.

### Experiment 2

Experimental setup is shown in Fig. 2A. We collected data from 26 subjects, but 2 were removed due to missing data for >= 15% of trials (final n=24, 22.42 ± 4.0 years old, 11 females). In this experiment, the perturbation was always +5°. Following a baseline period of 200 trials (80 straight vertical, 80 vertical oblique), adaptation blocks were alternated with error clamp blocks, as shown in Figs. 2B and 3A. The first experimental block was an adaptation block of 150 total trials (120 vertical trials), and the subsequent blocks were of 100 total trials (80 vertical trials) until the final error clamp block (150 total trials, 120 vertical trials). Total experiment length was 1700 trials (1280 vertical trials).

**Figure 2.**
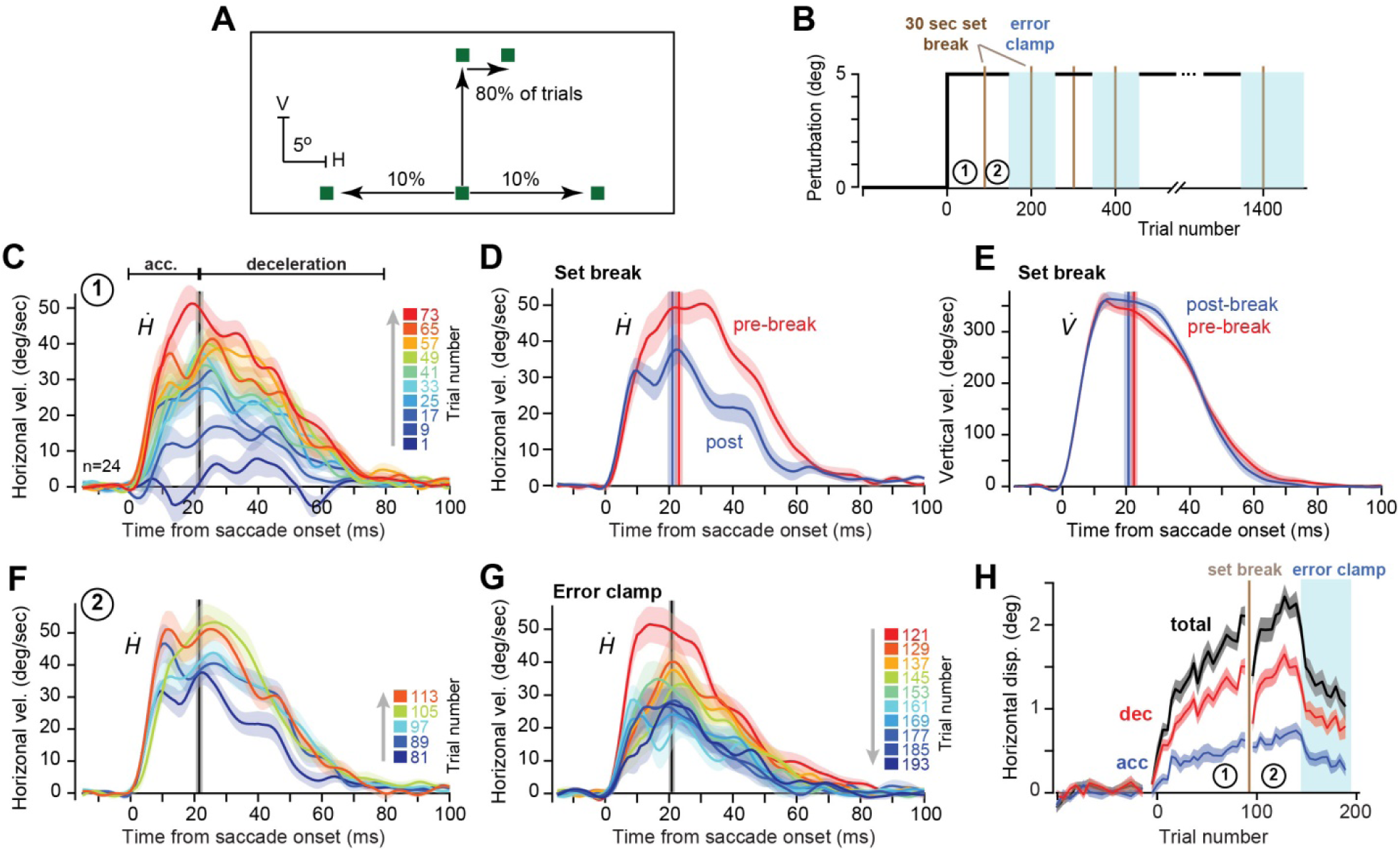
Acceleration and deceleration period commands adapt at different rates. **A**. Experimental paradigm. Following a vertical primary saccade, subjects experienced a rightward horizontal endpoint error. **B**. Training blocks (blocks 1 and 2 are labeled). **C**. Change in horizontal velocity during block 1 of the perturbation trials (data are within-subject change with respect to baseline). While one-trial learning exhibited corrective response only during deceleration, multi-trial learning exhibited corrective response during both acceleration and deceleration. **D**. Effect of set break on horizontal velocity. **E.** Effect of set break on vertical velocity. In contrast to the decay present in horizontal velocity, vertical velocity did not show decay. **F.** Re-learning in block 2. **G.** Decay of horizontal velocity during the error-clamp block. **H**. Total displacement due to the horizontal commands produced during the acceleration and deceleration periods. Deceleration period commands adapt at a faster rate during the perturbation block, but also exhibit greater loss during the set break. Both commands show decay during the error-clamp block. Error bars are between subject SEM.

**Figure 3.**
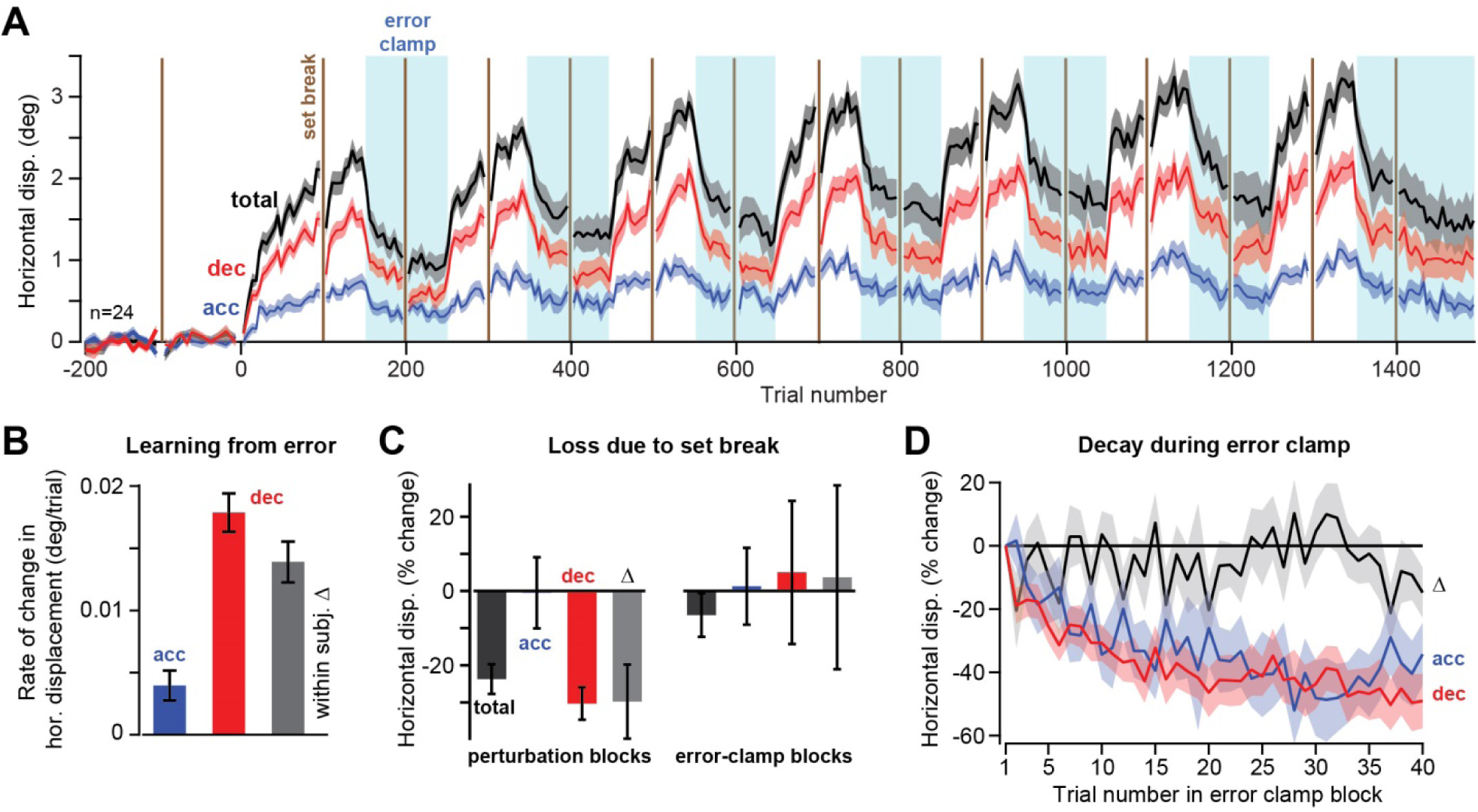
Acceleration and deceleration period commands exhibit some of the properties of the fast and slow system. **A**. Horizontal displacement produced by the acceleration and deceleration period commands during the various perturbation and error clamp blocks. **B**. Trial-to-trial rate of change in the horizontal displacement produced by the acceleration and deceleration period commands during the perturbation blocks. Rate of change was faster in the deceleration period. **C**. Effect of set-break on the horizontal displacement (within-subject percent change from before to after the set-break, bin size is 4 trials). Δ is the within subject difference between deceleration and acceleration. When the set-break was during the perturbation blocks, there was significant decay in the deceleration period commands, with little or no change in the acceleration period. In contrast, in the error-clamp blocks, both commands showed little or no decay. **D**. The acceleration and deceleration period commands decayed at similar rates during the error-clamp period. Error bars are between subject SEM.

In error clamp trials we used the current position, velocity, and acceleration of the eye when saccade speed fell below 100 degrees/sec during a saccade to predict where the saccade would end and placed the target at that location. We included a 30 second set break every 100 trials.

### Experiment 3

Experimental setup is shown in Fig. 5A. We collected data from n=40 subjects, 26.95 ± 4.86 years old, 10 females. Following a similar baseline period as in Experiment 2, subjects experienced a consistent 5° perturbation. As before, the perturbation direction was always perpendicular to the primary target direction. However, the subjects were divided into two equal groups in which the primary target was upward (vertical), or rightward (horizontal). The perturbation direction was maintained for 525 total trials (420 perturbed trials), after which it switched direction, as shown in Fig. 5A, lower subplot: that is, the initial perturbation trials were followed by a consistent 5° perturbation in the opposite direction for 50 total trials (40 perturbed trials). Subjects then finished the experiment with 125 total trials in error clamp (100 in the primary direction).

**Figure 4.**
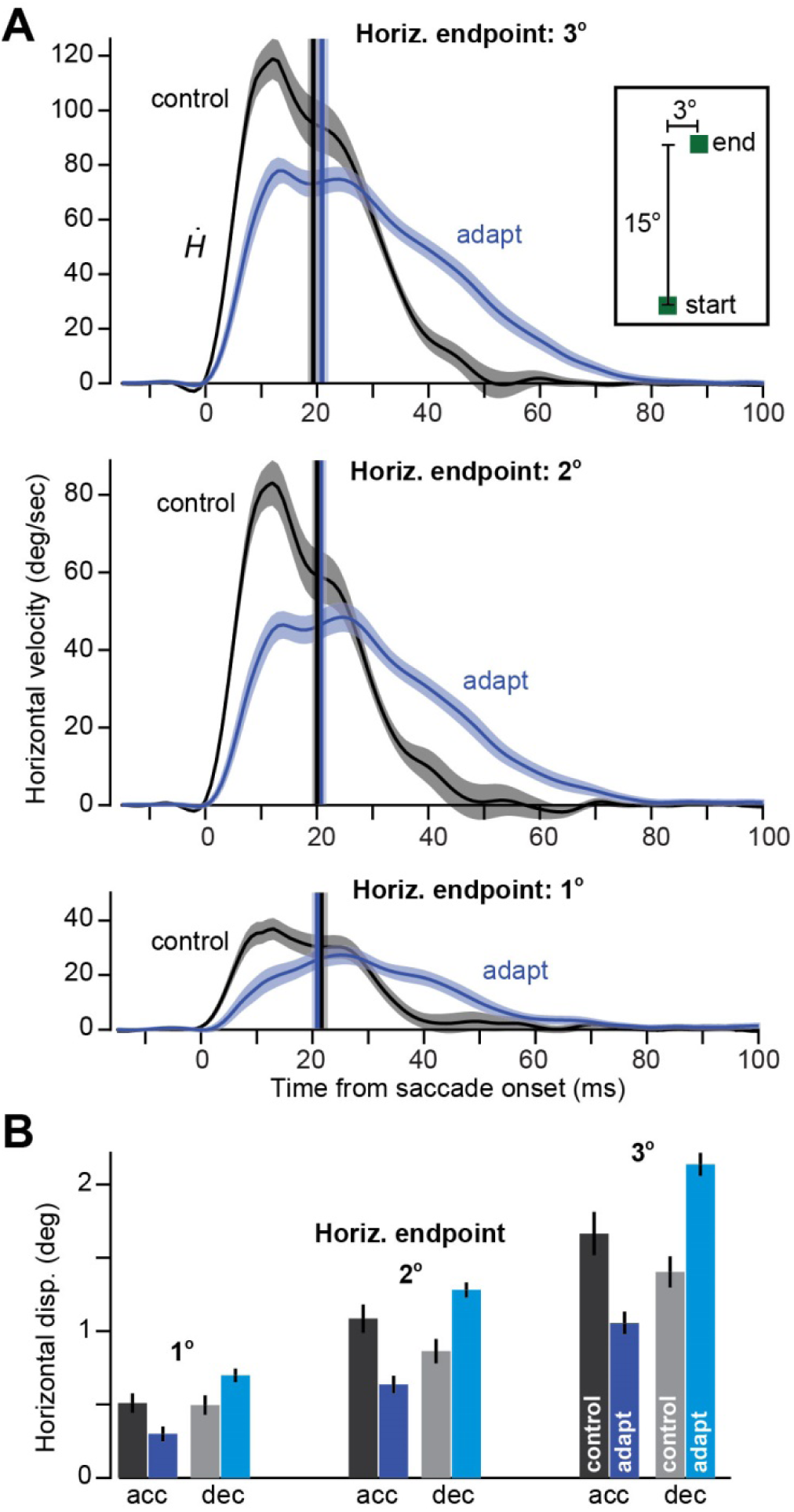
Adapted saccades arrive at a given endpoint with a trajectory that is quite different than control saccades. **A**. Horizontal velocity of control saccades and adapted saccades. In control trials, targets are presented at (+1°,15°), (+2°,15°), and (+3°,15°). In adapted saccades, the same endpoint is achieved but through learning. Adapted saccades exhibit smaller acceleration period horizontal displacement, but larger deceleration period horizontal displacement (with respect to baseline). **B**. A control saccade tends to have equal contributions to its horizontal displacement due to the acceleration and deceleration period commands. In comparison, an adapted saccade has most of its horizontal displacement due to the deceleration period commands. Bars are grouped based on size of endpoint horizontal displacement. Error bars are between subject SEM.

**Figure 5.**
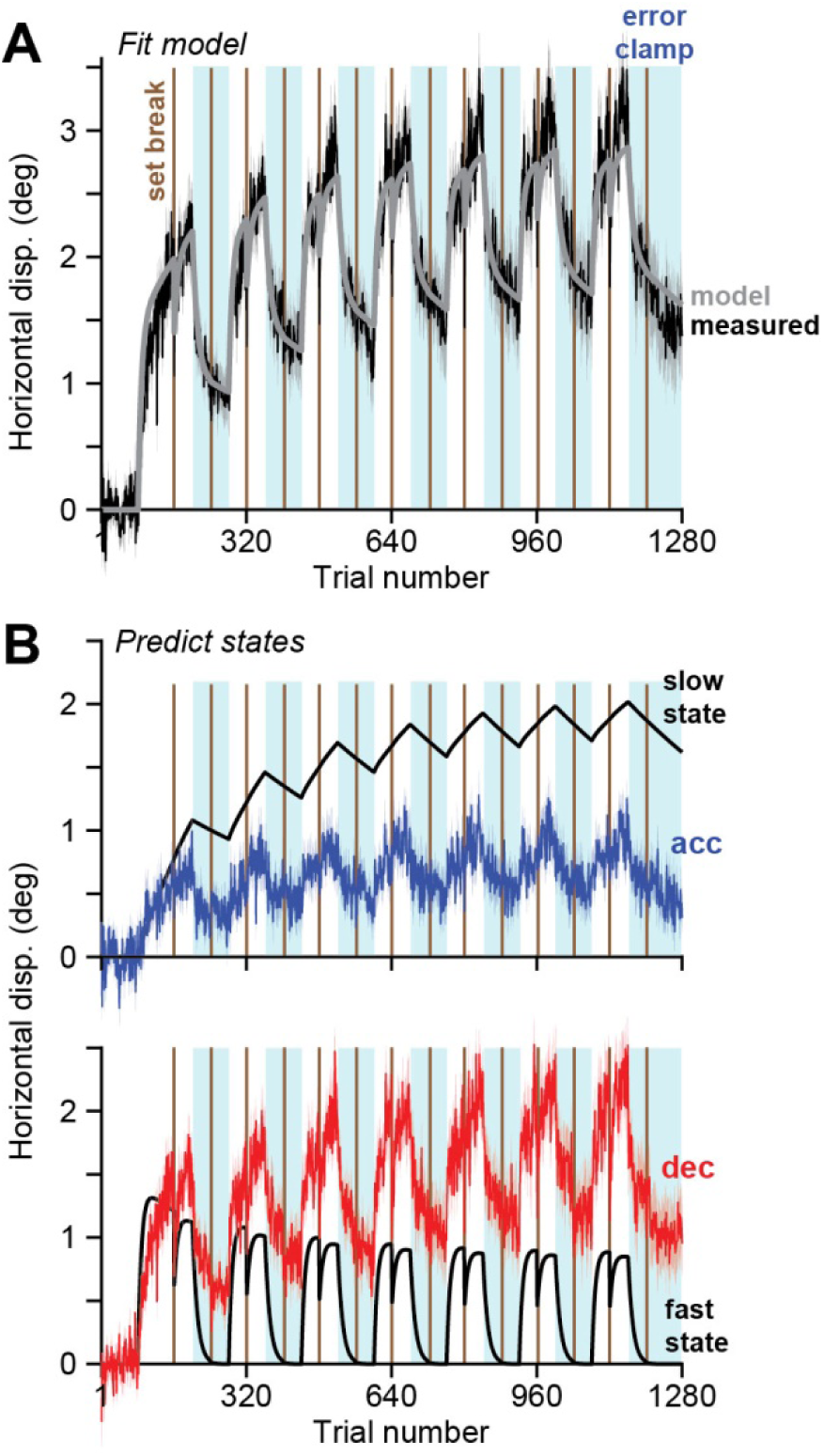
The 2-state model fits total behavior well, but does not predict the components of that behavior. **A**. Model fit to data (total horizontal displacement) in Exp. 2. **B**. The fast and slow states that are predicted by the model, and the underlying acceleration and deceleration period displacements. Note that in the first perturbation block, the predicted states are similar to the measured data. The measured data and predicts states diverge because in error clamp trials, the predicted states decay at different rates. The measured data, however, exhibits similar decay rates during the error clamp trials for the acceleration and deceleration period commands. Error bars are presented only for the measure data and are between subject SEM. Model was fit to the mean of the data.

### Data Analysis

All analyses were performed using in-house code written in MATLAB (The Mathworks, Natick, MA USA), except the repeated measures ANOVAs, which were performed in Rstudio for R. Position and velocity data from the eye were filtered off-line using a 3^rd^-order low-pass Butterworth filter with a cutoff frequency of 200 Hz prior to analysis.

Trial to trial analysis for Experiment 1 relied on the endpoint error that the subjects experienced on any given trial. We calculated error as the final position of the target minus the position of the eye at saccade end and defined errors less than −2.5° as negative errors (H-), and errors of greater than +2.5° as positive errors (H+). We then analyzed the change in horizontal position, displacement, and velocity, as well as the change in vertical velocity, from the trial in which the error was experienced to the next trial (Fig. 1E). To compute the time during the saccade in which the motor commands corrected for the previous error (Fig. 1G), we flipped the sign of the response to H- errors and combined them with the responses to H+ errors to obtain a single response to error profile for each subject. We then followed the procedures described by (Peel et al., 2017) to find the timing of the inflection point in the trajectory, resulting in the data shown in Fig. 1G.

For Experiments 2 and 3, we used a bin size of 4 trials and then performed a within-subject repeated measure ANOVA with bin and movement phase (acceleration and deceleration) as main effects including the interaction and horizontal displacement as the outcome measure. Learning rates were computed as the mean increase in horizontal displacement during acceleration and deceleration divided by the number of trials (Figs. 3B and 6B). Loss due to set break was computed as the average percent decay in each phase of the movement and the within-subject difference between the two movement phases (Fig. 3C) after applying a minimum threshold of an absolute value of at least 0.25° in each phase of the movement (2 subjects excluded, n=22). We performed t-tests for significant mean change for each movement phase and for difference in means between the two movement phases for both adaptation rate and loss. We performed a within-subject repeated measure ANOVA for the percent retention during the average error clamp block (within subject mean of 7 error clamp periods) with main effects of movement phase and trial as well as their interaction (Fig. 3D). As a control for Experiment 2, we compared the relative contributions of the acceleration and deceleration phases to total horizontal displacement for baseline (un-adapted) oblique saccades and adapted saccades with similar amounts of total horizontal displacement. We first performed a 2-way ANOVA with main effects of movement phase and condition (baseline vs. adapted) including the interaction term. We further calculated the percent contribution of acceleration and deceleration collapsed across the 3 levels of horizontal displacement (1°, 2°, and 3°) and performed paired two-tailed t-tests.

We quantified spontaneous recovery by using 3 key periods of 8 trials each, labeled in Fig. 5B, where *t*_1_ is the end of the initial adaptation period, *t*_2_ is the end of the de-adaptation period, and *t*_3_ is the peak of the spontaneous recovery curve (immediately after the final set break). We defined spontaneous recovery as the difference between displacements at *t*_3_ and *t*_2_ divided by the displacement at *t*_1_ (Fig. 5D), that is: [*x*(*t*_3_) − *x*(*t*_2_)] / *x*(*t*_1_), data in Fig. 6D.

**Figure 6.**
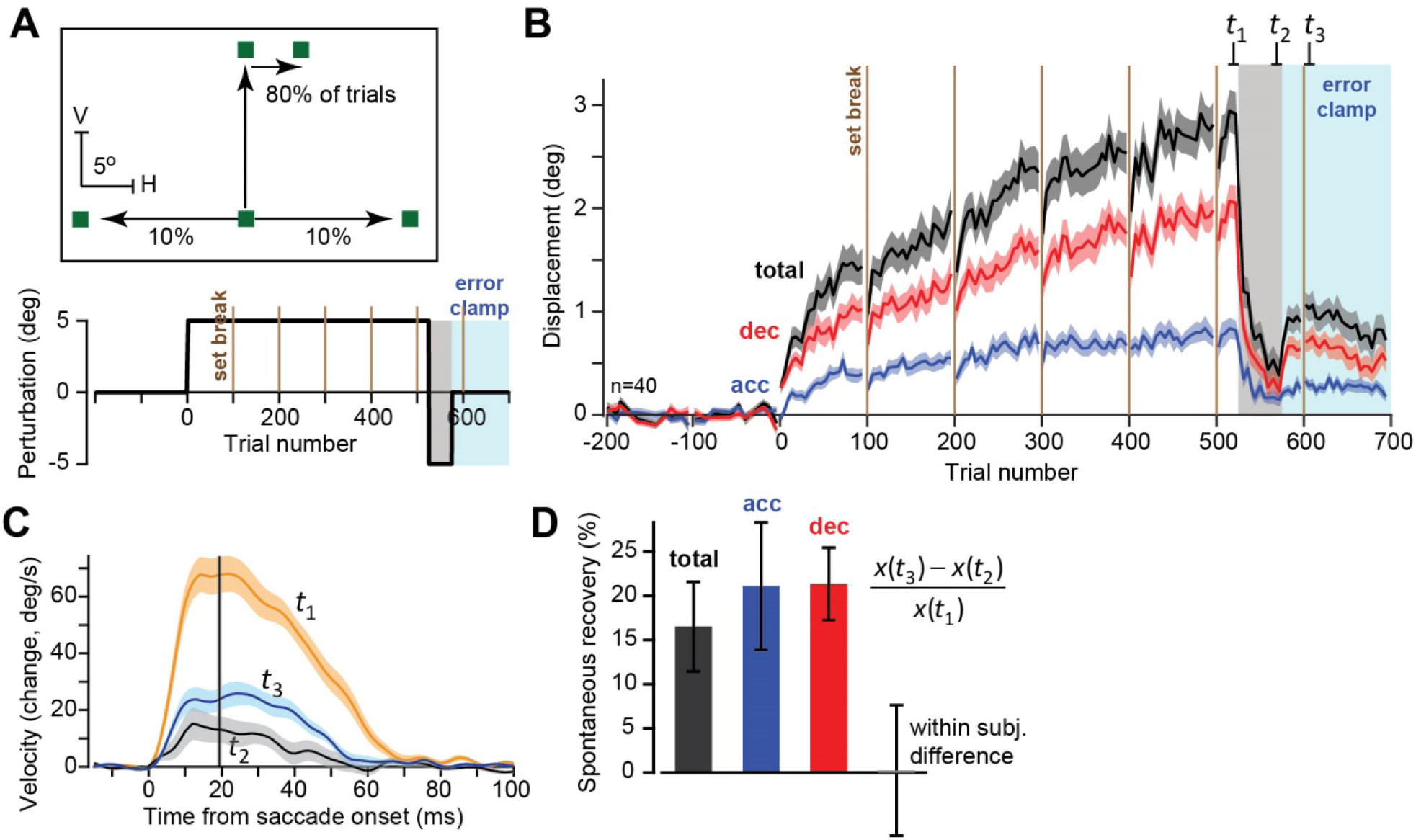
Experiment 3. Both acceleration and deceleration period commands exhibit spontaneous recovery. **A**. Experimental design. Perturbation block is followed by extinction, and concludes with a block of error-clamp trials. **B**. Saccade displacement in the direction of error during the acceleration and deceleration periods. Behavior in the error-clamp block exhibits spontaneous recovery. **C**. Eye velocity in the direction of error during the trials at end of adaptation (*t*_1_), at end of extinction (*t*_2_), and during error-clamp (*t*_3_). **D.** Spontaneous recovery, defined as [*x*(*t*_3_) − *x*(*t*_2_)] / *x*(*t*_1_).

### State-space models

After experience of a movement error, humans and other animals change their behavior on the subsequent trial. In the absence of error, adapted behavior decays over time. Here we used a state-space model to capture this process of error-based learning in our cross-axis saccade adaptation task (Donchin et al., 2003; Smith et al., 2006; Thoroughman & Shadmehr, 2000). The state-space model describes behavior as the summation of underlying hidden processes. That is, different brain regions, or neuronal circuits, each learn from error, and then combine their adapted states to produce the measured behavior. We represent each of these internal states via the vector ***x***.

Here we investigated the possibility that different parts of the same movement are supported by distinct internal states. Specifically, we focused on the acceleration and deceleration components of a single saccade. In Experiment 3 we found that both of these movement components undergo spontaneous recovery, suggesting that each are supported by at least two individual states of learning (Smith et al., 2006). Therefore, in total our state-space model requires four internal states, a slow state (*x*_*A,s*_) and fast state (*x*_*A,f*_) for acceleration, and a separate slow state (*x*_*D,s*_) and fast state (*x*_*D,f*_) for deceleration (Model 2, Fig. 7B). Our state vector, is a four-dimensional vector of the form ***x*** = [*x*_*A,s*_ *x*_*A,f*_ *x*_*D,s*_ *x*_*D,f*_]^*T*^. The internal states change from trial *n* to trial *n*+1 due to two different processes: learning from an error, e^(*n*)^, and trial-by-trial decay, according to Equation (1):

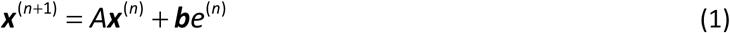

**Figure 7.**
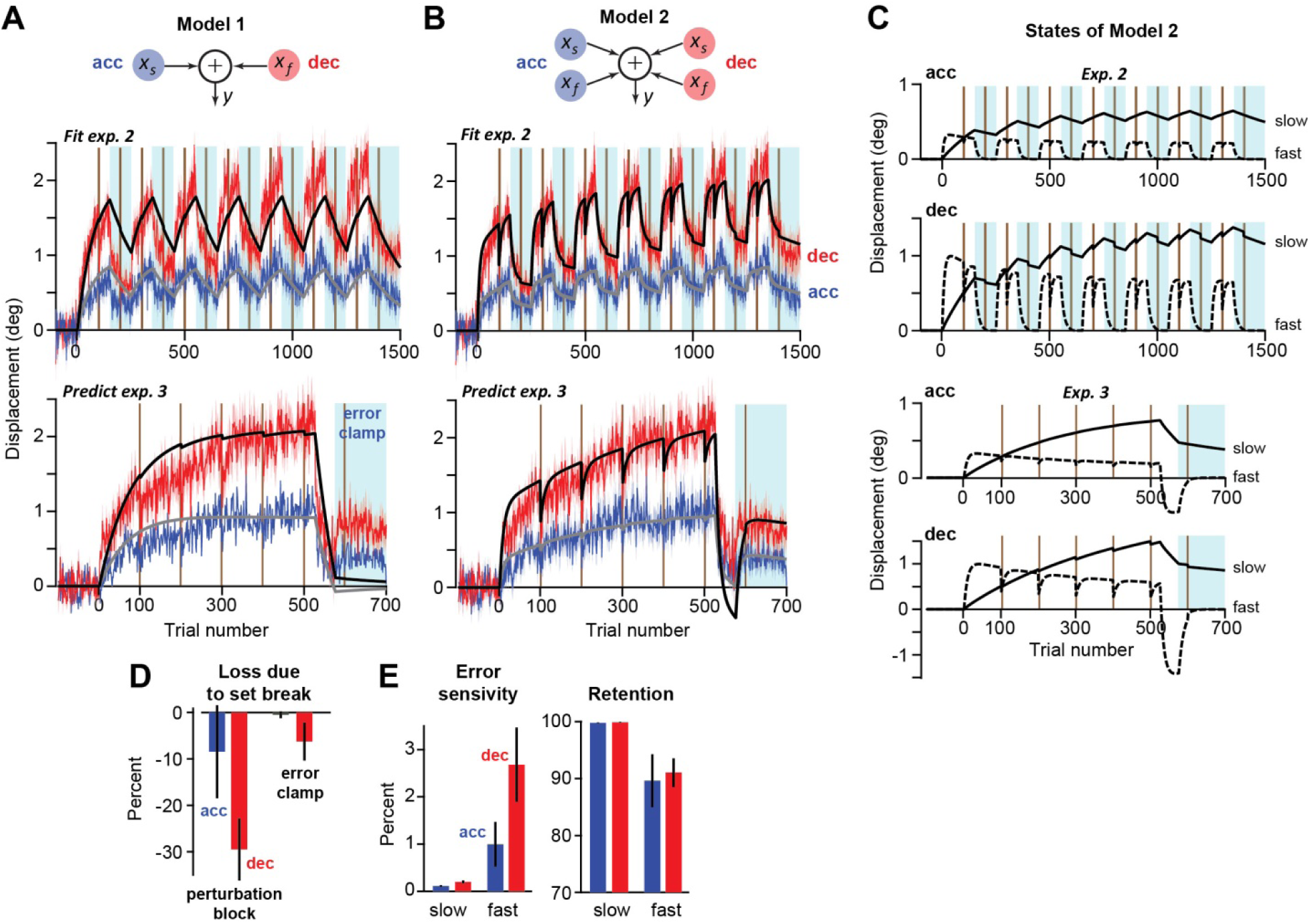
Two competing models of spontaneous recovery. In each case, the model was fit to the trial-by-trial acceleration and deceleration period commands in Exp. 2, and then used for predicting the measured data in Exp. 3. **A**. Model 1 has two states. Each state represents either the acceleration or the deceleration period commands. Each state learns from endpoint error and decays with time. The model fits the data in Exp. 2 well, and also predicts the perturbation period data in Exp. 3 reasonably well. However, the model does not correctly predict spontaneous recovery patterns during the error-clamp block of Exp. 3. **B**. Model 2 has four states: two states represent the acceleration period commands, and two states represent the deceleration period commands. This model correctly predicts much of the behavior in Exp. 3, including spontaneous recovery. **C**. States of Model 2 during Exp. 2 (top row), and Exp. 3 (bottom row). In Exp. 3, spontaneous recovery is present in both acceleration and deceleration period commands because both have a slow state that resists extinction. **D**. Model 2 correctly accounts for the fact that set breaks during the perturbation blocks produce a small loss in acceleration period commands, and a large loss in deceleration period commands (Experiment 2, compare to Fig. 3C). In contrast, during the error-clamp blocks, the same set-breaks produce much smaller losses to these commands. This is because during error-clamp blocks, the fast states in each type of command have largely decayed, resulting in resistance to loss in set-breaks. **E**. Parameters of Model 2. Error bars are standard deviation, estimated via boot-strap.

Forgetting is controlled by the retention matrix *A*, which enforces that each state decays exponentially over trials, in an independent manner. Therefore, *A* is a diagonal matrix whose main diagonal entries are the individual retention factors for the slow and fast acceleration and deceleration states: *a*_*A*,s_, *a*_*A,f*_, *a*_*D,s*_, and *a*_*D,f*_. Next, the rate of learning is controlled by the error sensitivity vector ***b***. The error sensitivity vector consists of the individual error sensitivities for each internal state: *b*_*A*,s_, *b*_*A,f*_, *b*_*D,s*_, and *b*_*D,f*_. The two-state model posits that slow internal states learn more slowly from the experience of error than fast internal states, but forget less from one trial to the next. These conditions are enforced by the following four inequalities:

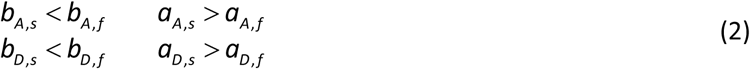

While we cannot directly measure the internal states of an individual, we can measure their combined effect on the adapted movement:

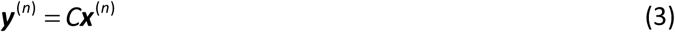

Here, the movement vector ***y***, consists of an acceleration component *y*_*A*_ and a deceleration component *y*_*D*_. Therefore, the matrix *C* takes the form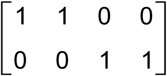, ensuring that the slow and fast acceleration states contribute to the acceleration component of the movement, and the slow and fast deceleration states contribute to the deceleration component of the movement.

Our experiments consisted of a combination of trial conditions. On some trials, a perturbation *r* was applied to the target position. On these trials, the error experienced by the participant is equal to the difference between the perturbation, and the total displacement of the eye, *y*_*A*_ + *y*_*D*_. On error-clamp trials, movement errors were entirely eliminated by moving the target to the terminal eye position.

Therefore, the error in Eq. (1) has two possible forms:

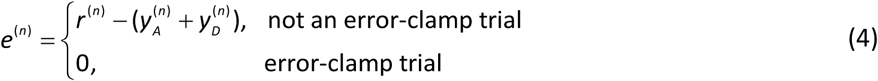

Lastly, after some trials, participants were allowed to rest during a set break. Here we modeled the decay that accompanies the passage of time during a set break (Scott T. Albert & Shadmehr, 2018) by multiplying the internal states on the right-hand-side of Eq. (1) by a decay matrix *D*:

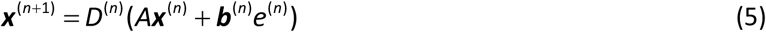

We allowed decay to differ for each of the four internal states. Therefore, the decay matrix consisted of a diagonal matrix with main diagonal entries *d*_*A,s*_, *d*_*A,f*_, *d*_*D,s*_, and *d*_*D,f*_. On normal trials not followed by a set break, the *D* matrix was set equal to the identity matrix. In total, our state-space model is represented by the system:

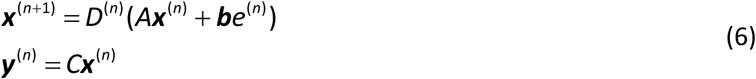

where the error *e* is given by Eq. (4), and inequalities describing retention factors and error sensitivities are given by Eq. (2).

To illustrate that slow and fast states of learning are required to explain the acceleration component and deceleration component of the measured behavior, we compared our state-space model in Eq. (6) with a simpler version consisting of only two states (Model 1, Fig. 7A). In other words, we assumed that both acceleration and deceleration each consisted of a single state.

This reduced model has the same structure as the state-space model in Eq. (6), with the following updated terms. *A* is a 2 × 2 diagonal matrix with a retention factor for acceleration (*a*_*A*_) and deceleration (*a*_*D*_) along the main diagonal. The error sensitivity vector ***b*** has only two components, an error sensitivity for acceleration (*b*_*A*_) and another for deceleration (*b*_*D*_). The *C* matrix is in this case a 2 × 2 identity matrix. Finally, the decay matrix for set breaks is a 2 x 2 diagonal matrix with entries *d*_*A*_ and *d*_*D*_. Unlike the constraints in Eq. (2), we did not constrain this simpler model with any inequalities restricting the relationship among model parameters. In total, Model 1 had six parameters, *a*_*A*_, *a*_*D*_, *b*_*A*_, *b*_*D*_, *d*_*A*_, and *d*_*D*_. In comparison, Model 2 required twelve parameters: *a*_*A,s*_, *a*_*A,f*_, *a*_*D,s*_, *a*_*D,f*_, *b*_*A,s*_, *b*_*A,f*_, *b*_*D,s*_, *b*_*D,f*_, *d*_*A,s*_, *d*_*A,f*_, *d*_*D,s*,_ and *d*_*D,f*_.

### Fitting state-space model to measured behavior

We fit our state-space models to the behavior recorded in Experiment 2, and then used the resulting models to predict behavior in Experiment 3. There were six parameters for Model 1 (Fig. 7A), and twelve parameters for Model 2 (Fig. 7B). We fit each model to the trial-by-trial sequence of the mean population data, and used boot-strapping techniques to estimate the variance of the model parameters. Once the model parameters were identified from data in Experiment 2, we used it to predict behavior in Experiment 3.

To fit the state-space model, we used a least-squares approach. Least-squares approaches are susceptible to poor fit quality if not carefully constrained (Scott T. Albert & Shadmehr, 2018). Here, we ensured excellent fit quality, by constraining the initial internal states to zero, and by using paradigms that have a combination of many conditions (perturbations of different signs, error-clamp blocks, and set breaks). In these cases, least-squares techniques have been shown to perform similarly to other maximum likelihood techniques (Albert & Shadmehr, 2018).

We fit the 6-parameter (θ_6_) and 12-parameter (θ_12_) models to the mean behavior recorded in Experiment 2 by minimizing the following squared error cost function:

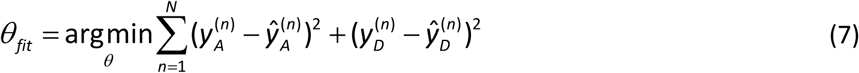

This minimization was performed using *fmincon* in MATLAB R2018a. Minimization of Eq. (7) was subject to the linear inequality constraints in Eq. (2) for the 12-parameter fit, but not for the 6-parameter fit. Finally, all parameters were bounded between 0 and 1. To ensure that the global minimum of Eq. (7) was identified, we performed 100 iterations of *fmincon*, each with a different randomly specified initial condition. The parameter set associated with the smallest squared error was selected. We then used this parameter set to simulate the expected performance in Experiment 3.

### Canonical two-state model of behavior

In a separate analysis, we considered the possibility that the canonical fast and slow states of adaptation (Smith et al., 2006) corresponded to the deceleration and acceleration components of the movement (Fig. 5). To test this, we fit the standard two-state model of adaptation (Albert and Shadmehr, 2018) to the measured behavior along the horizontal axis, ignorant of the acceleration and deceleration components of the saccade. We identified the parameter set that minimized the sum of squared error between the model prediction and mean behavior in Experiment 2:

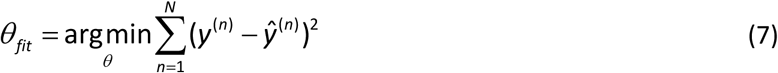

where θ consisted of a slow state with retention factor *a*_*s*_, error sensitivity *b*_*s*_, and time decay factor *d*_*s*_, and a fast state with retention factor *a*_*f*_, error sensitivity *b*_*f*_, and time decay factor *d*_*f*_. As in Eq. (2), we required the fast state to have a greater error sensitivity, but smaller retention factor than the slow state of adaptation.

## Results

We asked healthy human participants (n=101 in total) to make a saccade toward a visual target at 15°. As the primary saccade started, we moved the target perpendicular to the direction of the target by 5°, thus presenting a visual error that was followed by a corrective saccade. The experience of error produced a small but robust change on the subsequent primary saccade toward the 15° target. We quantified the trial-to-trial change in saccade trajectory, which unmasked differing properties of adaptation in the acceleration and deceleration periods.

### Single trial learning: learning from error differed during the acceleration and deceleration periods of the movement

In Exp. 1, participants (n=37) made primary saccades mainly in the vertical direction (80% of the trials) and occasionally experienced an error (Fig. 1A): during the primary saccade the target was displaced horizontally to the right or left (13% of trials in each case, Fig. 1B), resulting in a visual error. The direction of error was random but balanced between right and left. To measure learning from error, we considered pairs of trials, *n* and *n* + 1, in which the primary saccade in both trials was to the vertical target, and the error occurred in trial *n*. Learning from error was measured as the trial-to-trial change in the horizontal and vertical eye velocity (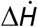 and 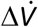) of the primary saccade, that is: 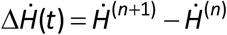.

Fig. 1C displays the eye velocity for all primary vertical saccades, and Fig. 1D presents the trial-to-trial change following experience of a positive *H*^+^ or a negative *H*^−^ error in trial *n*. The vertical gray line in Fig. 1D marks the time of peak saccade velocity, defined as 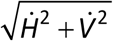. Thus, the period before the peak velocity is the acceleration period, and the period after is the deceleration period.

We had expected that experience of the horizontal error would produce a simple adaptive change in the subsequent trial: a small increase in the horizontal velocity in the direction that would reduce the error, along with little or no change in the vertical velocity. As expected, there was no error-dependent change in the vertical velocity (left panel of Fig. 1D): in the deceleration period, there was a slight increase in vertical velocity, but the effect did not differ between *H*^+^ and *H*^−^ errors (5.94 ± 0.77 vs. 5.27 ± 0.57 deg/sec, respectively, paired t-test for difference in means p=0.42). Surprisingly, we found that the change in horizontal velocity was bimodal and differed between acceleration and deceleration. For example, following a positive error *H*^+^, during the acceleration of the primary saccade 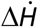 was negative, thus pulling the eyes in the wrong direction with respect to the error (green trace, right panel of Fig. 1D). However, during the deceleration period 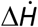 became positive, pulling the eyes in the direction that compensated for the error. A similar bimodal pattern was present following experience of *H*^−^ error (brown curve, right panel of Fig. 1D).

The effect of this one-trial learning on eye trajectory is displayed in Fig. 1E: experience of an error to the right (*H*^+^, green trace, Fig. 1E) was followed by a primary saccade that during its acceleration changed the eye trajectory in the wrong direction. However, as the saccade continued into the deceleration phase, the eye was steered correctly, thus partially compensating for the error experienced in the previous trial. As a result, following an error, the next saccade improved in the sense that its endpoint exhibited a smaller error (had the perturbation repeated). However, this reduced endpoint error was due to commands that arrived primarily during the deceleration period.

To better quantify the exact time when the commands correctly changed eye trajectory, we analyzed data of each participant separately and looked for the onset time in which 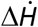 corrected the eye trajectory in the direction of the previous error. The results revealed that error compensation did not begin at saccade initiation (Fig. 1G, left subplot, mean 24.14 ± 1.12 ms, t(36) = 21.54, p = 7.4×10^−22^). Rather, the compensation started 3.67 ± 1.05 ms following peak velocity, i.e., almost precisely at deceleration onset (right subplot, Fig. 1G).

In summary, following experience of an endpoint error, the subsequent primary saccade included small, corrective commands that steered the eyes, thus reducing the error. However, the expression of these corrective commands differed between the acceleration and the deceleration periods. Only the deceleration period commands corrected the eye’s trajectory, reducing the endpoint error. If we assume that learning that follows experience of a single error is due to the hypothesized fast adaptive process, then the results of Exp. 1 imply that the fast process improved behavior mainly through changes in commands that were expressed during deceleration, not acceleration.

### Adaptation of deceleration period commands exhibited high sensitivity to error, but poor retention

We wondered whether the response to error in the deceleration period would remain dissociated if errors were not random, but always in the same direction. In Exp. 2, participants (n=24) made primary saccades mainly in the vertical direction (80% of the trials) and always experienced a horizontal error to the right (Fig. 2A). In addition to the perturbation trials, we also included set breaks and error clamp trials (Fig. 2B).

During the first perturbation block (Fig. 2B, part 1), horizontal velocity gradually increased from baseline (Fig. 2C). Our naïve expectation was that after a few trials of error experience, horizontal velocity will be a scaled version of the response that we had recorded following a single trial. However, whereas single trial learning had exhibited a bimodal response, multi-trial learning produced a unimodal response that was entirely in the direction of error.

We quantified the effects of the acceleration and deceleration period commands by integrating the horizontal velocity during each period (Fig. 2H). The commands during both periods moved the eyes in the direction of error, but the displacement arising from the deceleration period appeared to grow at a faster rate. We tested this observation with a repeated measures ANOVA with trial (bin of 4) and movement phase (acceleration or deceleration) as main factors. As expected, the main effect of trial was highly significant (F(9,207) = 51.97, p = 2×10^−16^), reflecting robust adaptation, and the main effect of movement phase was also significant (F(1,23) = 69.18, p = 2.19×10^−8^), indicating an offset between acceleration and deceleration. Crucially, the interaction of trial and movement phase was highly significant (F(9,207) = 6.78, p = 1.64×10^−8^), demonstrating that the rate of adaption was different between acceleration and deceleration. We confirmed that adaptation during the deceleration phase increased at a faster rate with a paired t-test(Fig. 2H, 0.017 ± 0.001 deg/trial vs. 0.008 ± 0.002 deg/trial, t(23) = 3.80, p = 0.001). That is, during the perturbation block, the trial-to-trial rate of change was faster for the commands that arrived in the deceleration period as compared to the acceleration period.

Following the set break, there was forgetting: horizontal velocity decreased toward baseline (Fig. 2D). The effect was somewhat larger in the deceleration period (Fig. 2H, set break, percent change, −9.5 ± 5% for acceleration period, −35.8 ± 5% for deceleration period). In contrast, the effect of set break was negligible in the vertical velocity (vertical velocity slightly increased following the set break, Fig. 2E). Thus, the set break induced forgetting, but the loss was not uniformly expressed throughout the movement: it appeared to be larger for the commands during the deceleration period.

In the subsequent re-exposure to the perturbation (Fig. 2F), horizontal velocity increased. Again, the rate of change in the deceleration period was larger than the rate of change in the acceleration period (Fig. 2H, label 2, trial X movement phase interaction F(4,92) = 5,48, p = 0.0005, rates 0.016 ± 0.003 deg/trial vs. 0.006 ± 0.002 deg/trial, t(23) = 2.48, p = 0.02). Finally, in the error clamp block that ensued, horizontal velocity exhibited a gradual decay (Fig. 2G).

These patterns repeated in the subsequent blocks of training (Fig. 3A). During the perturbation blocks, the deceleration period commands adapted at a faster rate than the acceleration period commands (Fig. 3B, trial X movement phase interaction F(4,92) = 13.02, p = 1.92×10^−8^, rate within subject difference 0.019 ± 0.003 deg/trial, t(23) = 6.92, p = 4.72×10^−7^). Following a set break, the deceleration period commands suffered a substantial loss (Fig. 3C, left subplot, −30 ± 4%, t(21) = −6.95, p = 7.24×10^−7^). In contrast, there was little or no loss in the acceleration period commands (Fig 3C, left subplot, −0.5 ± 9.6%, t(21) = −0.05), p = 0.96). Both the acceleration and deceleration period commands decayed during the error clamp blocks (Fig. 3D, RMANOVA main effect of bin F(39, 897) = 8.12, p = 2×10^− 16^). However, the rates of decay were indistinguishable (main effect of movement phase F(1,23) = 0.686, p = 0.42, trial X movement phase interaction F(39,897) = 1.27, p = 0.13, rates 0.68 ± 0.31%/trial vs. 1.17 ± 0.21%/trial, test of within subject difference t(23) = −1.53), p = 0.14).

In summary, in response to repeated endpoint errors, the primary saccade gradually adapted: the brain learned to augment the vertical commands with horizontal commands. Because single-trial learning had demonstrated that the response to error differed between the acceleration and deceleration periods, we analyzed adaptation by separately integrating the horizontal velocity in these two periods and found that the deceleration period commands adapted rapidly, but suffered from a large loss during set breaks. In contrast, the acceleration period commands adapted slowly, and exhibited little or no loss during the same set breaks. During the error clamp block, the commands in both acceleration and deceleration periods exhibited decay, but there were no differences in the rates of decay.

### Set breaks induced forgetting, but not during the error-clamp block

Behavior following set breaks presented an interesting puzzle. The set breaks in the perturbation blocks induced partial loss of the learned behavior (Fig. 3C, left subplot): total horizontal displacement declined by 22 ± 3.8% (t(23) = −5.78, p = 6.92×10^−6^). This loss was largely due to reductions in the deceleration period commands (within subject difference between loss in the acceleration period vs. deceleration period, t(21) = −2.99, p = 0.007). However, when the same set breaks were given in the error clamp blocks, there was no significant change in behavior (within subject change in total horizontal displacement, −9.6 ± 6.7%, t(23) = −1.44, p = 0.16, Fig. 3C, right subplot). Indeed, set breaks during the perturbation blocks induced greater loss than set breaks during the error clamp blocks (within subject difference, 12.4 ± 5.8%, t(23) = 2.16, p = 0.042).

To summarize, there were two puzzling aspects of behavior: whereas exposure to error produced rapid adaptation of the deceleration period motor commands, removal of error (in the error-clamp block) produced comparable decay in the acceleration and deceleration periods. Whereas set breaks during perturbation blocks produced a large loss in the deceleration period commands, the same set breaks during the error clamp block produced little or no loss. We will consider these results shortly when we attempt to build a model.

### Control trials: deceleration period commands were specific to adapted saccades

By the end of training, in response to a vertical target at (0°,15°), the brain generated a saccade that moved the eyes to (3°,15°): the horizontal commands steered the eyes by roughly 3° as the eyes traveled 15° along the vertical direction (Fig. 3A). As noted, this 3° change in horizontal displacement was more strongly influenced by the deceleration component of the saccade, which responded more vigorously to the occurrence of error (Figs. 1, 2, and 3). Could this disparity between the decelerative and accelerative contributions to adaptation be trivially related to the relative durations of eye acceleration and deceleration? That is, the deceleration component lasted approximately 40 ms, allowing nearly twice the amount of time for correction than that of the acceleration component which lasted approximately 20 ms (Figs. 1 and 2). To consider this possibility, we compared the trajectory of adapted saccades to oblique saccades that had the same endpoint, but were made in a control condition in which the primary target was presented at (3°,15°).

In the control block of trials, before introduction of perturbations, we included trials in which the target was presented at (1°,15°), (2°,15°), …, (5°,15°). Fig. 4A plots the horizontal velocity in the control and adapted conditions. The saccades in these two conditions differed markedly. For example, the top plot of Fig. 4A displays horizontal velocities for saccades that ended with a 3° horizontal displacement. In the control condition, the acceleration and deceleration period commands equally contributed to the horizontal displacement. In contrast, in the adapted condition, the 3° endpoint displacement was achieved mainly via commands that arrived in the deceleration period (within-subject 2-way ANOVA, main effect of adaptation vs. baseline F(1,23) = 5.80, p = 0.02, main effect of acceleration vs. deceleration F(1,23) = 1.33, p = 0.26, interaction F(1,23) = 49.55, p = 3.58×10^−7^). This pattern repeated across the various control and adapted saccades (Fig. 4B, within subject difference between percent contribution of acceleration and deceleration commands collapsed across 1, 2, and 3 deg endpoints, baseline vs. adapt, t(23) = 5.13, p = 3.42×10^−5^).

Therefore, in all saccades the deceleration period lasted longer than the acceleration period. However, in control saccades the horizontal displacement was due to roughly equal contributions from acceleration and deceleration period commands (in Fig. 4B, gray bars, deceleration minus acceleration, - 10 ± 11%, t(23) = −0.91, p = 0.37). In contrast, for adapted saccades a different pattern emerged. Even though the oculomotor learning system could have accomplished the desired horizontal displacement using acceleration and deceleration equally, it somehow was limited to express its contributions mainly during the deceleration period (blue bars in Fig. 4B, dec – acc: 34 ± 5%, t(23) = 7.12, p = 2.98×10^−7^).

### The standard model failed to predict the patterns of commands during adaptation

Acceleration period commands adapted slowly and exhibited little or no loss during set breaks (Fig. 3C). Deceleration period commands adapted rapidly and exhibited large loss during perturbation block set breaks. These dynamics were reminiscent of the slow and fast states of the 2-state model. However, there were also puzzling aspects of behavior: during the error clamp block, acceleration and deceleration period commands decayed at similar rates (Fig. 3D), and the set breaks produced little or no loss. Is there a way to reconcile all of these findings within a single framework?

We began by fitting the total horizontal displacement in Exp. 2 (total, Fig. 3A) to the standard 2- state model and found that it fit the data exceedingly well (Fig. 5A). However, the resulting fast and slow states bore little resemblance to the dynamics of the acceleration and deceleration period commands (Fig. 5B). In particular, the actual acceleration period commands exhibited a much smaller increase than that predicted by the slow state, and the actual deceleration period commands exhibited a much greater increase than that predicted by the fast state.

The mismatch between the model’s predictions and measured behavior was particularly evident in the error-clamp blocks. The model fit the total horizontal displacement quite well during the error clamp blocks (Fig. 5A), but predicted that the fast state would decay more rapidly than the slow state (black lines, Fig. 5B). However, the acceleration and deceleration commands decayed at similar rates in the error clamp blocks (Fig. 3D). As a result, the difference between model predictions and the measured behavior grew as the error clamp blocks repeated (Fig. 5B).

Thus, while the 2-state model could fit general behavior quite well (total endpoint displacements), its predictions regarding the underlying components of behavior (acceleration and deceleration period displacements) were incompatible with the measurements.

### Both the acceleration and deceleration period commands exhibited spontaneous recovery

Perhaps the most informative paradigm that helps expose the multiple timescales of motor memory is one in which learning is followed by extinction, thus resulting in spontaneous recovery. In Experiment 3, subjects (n=40) made primary saccades to a target at 15°, and then experienced an error perpendicular to the direction of the target (Fig. 6A). The design of the experiment was similar to Exp. 2, except that following adaptation, the direction of the perturbation reversed, thus encouraging extinction. Exp. 3 concluded with a block of error-clamp trials.

As before, we observed that the deceleration period commands adapted more rapidly than the acceleration period commands (Fig. 6B, RMANOVA main effect of bin F(26,1014) = 39.44, p = 2×10^−16^, main effect of movement phase F(1,39) = 97.84, p = 3.48×10^−12^, interaction F(26,1014) = 39.44, p = 7.80×10^−16^, rate of change, 0.005 ± 0.0005 deg/trial vs. 0.002 ± 0.0003 deg/trial, test of within subject difference t(39) = 6.19, p = 2.86×10^−7^). After 420 trials, saccade endpoint had changed by roughly 3° (trials labeled *t*_1_, Fig. 6B). To induce extinction, we reversed the direction of the visual error. This resulted in sharp reductions in the acceleration and deceleration period commands (gray region of Fig. 6B). By the end of the extinction trials, endpoint displacement was less than 0.5° (*t*_2_, Fig. 6B).

Following the extinction trials, we presented the participants with a block of error clamp trials. This induced spontaneous recovery: saccade endpoint spontaneously increased from 0.5° to around 1° (from *t*_2_ to *t*_3_, black line, Fig. 6B). We quantified spontaneous recovery as the change in the displacement from the end of extinction trials (time *t*_2_) to the error clamp block (time *t*_3_), that is: (*H* (*t*_3_) − *H* (*t*_2_)) / *H* (*t*_1_). This measure indicated that saccade endpoint exhibited roughly 16.5% spontaneous recovery (bar plot labeled “total”, Fig. 6D, t(34) = 3.26, p = 0.002). Of importance is the fact that spontaneous recovery was present both in the acceleration and the deceleration periods (Fig. 6D, 21 ± 7%, t(34) = 2.93, p=0.006 and 21 ± 4%, t(34) = 5.19, p=9.87×10^−6^). Indeed, spontaneous recovery did not differ between the two periods of the movement (within subject difference, 0.24 ± 7.38%, t(34) = 0.032, p = 0.97).

The patterns of spontaneous recovery in the deceleration period are noteworthy because they are opposite of what would be expected by the 2-state model. The deceleration period commands had exhibited rapid learning along with large forgetting (during set breaks), making them seem like a proxy for the fast process. According to the 2-state model, during the error-clamp trials all states should simply decay toward zero. In sharp contrast, both the deceleration as well as the acceleration period commands exhibited spontaneous recovery.

Behavior around the set breaks exhibited an interesting pattern. Whereas during the perturbation blocks the set breaks consistently produced forgetting, the set break during the error-clamp block did not produce forgetting (Fig. 6B, +0.17 ± 0.09°, t(39) = 1.79, p=0.08). This finding repeated what we had seen in Experiment 2: set breaks in one context (perturbation block) accompanied forgetting, but not in another context (error-clamp trials).

In summary, following extinction, there was spontaneous recovery of saccade endpoints toward the initially learned behavior. However, this was not because there were differential decay patterns in the acceleration and deceleration commands. Rather, commands during both periods exhibited spontaneous recovery. Finally, set breaks during the perturbation blocks induced forgetting. However, forgetting did not occur when the same set break was given during the error clamp block.

### A new model of spontaneous recovery

Is there a mathematical model that can in principle reproduce the rich body of data in the various experiments? We considered two alternatives (Fig. 7). In each case, we imagined that the acceleration and deceleration period commands were supported by distinct adaptive controllers. Each controller learned from error, but could express that learning only during a specific period of the movement. We fit each model to data in Experiment 2, and tested them by comparing their predictions to the measured data in Experiment 3.

Model 1 was a modified version of the standard 2-state system: acceleration and deceleration commands were served by distinct controllers, each supported by a single state that learned from error (Fig. 7A). Thus, in this model there were two states: both states changed following experience of error, and decayed with passage of time.

In Model 2, acceleration and deceleration were served by distinct controllers, but each was served by two states, fast and slow (Fig. 7B). Thus, in this model there were four states, with two states contributing to the acceleration phase, and two states contributing to the deceleration phase.

We found that both models fit the data in Exp. 2 quite well (acceleration and deceleration period displacements, Fig. 7A and 7B, top row). The resulting model parameter values for Model 1 were {*a*_*A*_=0.9923, *a*_*D*_=0.9938, *b*_*A*_=0.0034, *b*_*D*_=0.0066, *d*_*A*_=1.0, *d*_*D*_=0.9749} and Model 2 were {*a*_*A,s*_=0.9978, *a*_*D,s*_=0.9989, *a*_*A,f*_=0.8963, *a*_*D,f*_=0.9105, *b*_*A,s*_=0.0011, *b*_*D,s*_=0.002, *b*_*A,f*_=0.0099, *b*_*D,f*_=0.0268, *d*_*A,s*_=1.0, *d*_*D,s*_=0.9541, *d*_*A,f*_=0.763, *d*_*D,f*_*=*0.4303} (Model 2 parameters are plotted in Fig. 7E). Having found the parameters of each model, we used the resulting equations to make predictions for Experiment 3.

The second rows of Figs. 7A and 7B show the predictions of each model. In the perturbation and extinction blocks of Exp. 3, output of both models resembled the measured data (Fig. 7A and 7B, second row). However, Model 1 did not correctly predict dynamics around the set breaks. Furthermore, following extinction and during the subsequent error clamp period, Model 1 did not correctly predict the spontaneous recovery that was present in the acceleration and deceleration commands.

In comparison, Model 2 correctly predicted behavior at set breaks during the perturbation block, as well as the patterns of spontaneous recovery. The reason that Model 2 performed well can be gleaned by examining the evolution of its various states (Fig. 7C). Because the acceleration and deceleration period controllers were endowed with separate fast and slow states, both components of behavior exhibited spontaneous recovery. These two distinct states for each controller also allowed Model 2 to account for dynamics of behavior around the set breaks.

Recall that a key puzzle in our data was the effect of set breaks: when the set breaks occurred during the perturbation block, acceleration period commands did not decay (Fig. 3C), whereas deceleration period commands showed substantial decay. In contrast, when the same set breaks occurred during the error-clamp block, neither acceleration nor deceleration period commands exhibited decay (Fig. 3C). Remarkably, Model 2 reproduced both of these behaviors, as shown in Fig. 7B (top row), and quantified in Fig. 7D. Model 2 accounted for this diversity as follows: when the set breaks occurred during the perturbation block, the fast states were non-zero, and thus exhibited decay (Fig. 7C). However, when the set breaks occurred during the error-clamp block, the fast states had already decayed to zero, and thus could not decay any further. As a result, set breaks during perturbation blocks produced decay, but not during the error-clamp blocks.

Thus, by assuming that different controllers contributed to the motor commands in the acceleration and deceleration periods, and that each controller was supported by a fast and slow process, Model 2 was able to account for spontaneous recovery, as well as the rich patterns that were present in the acceleration and deceleration period commands.

## Discussion

In numerous paradigms, learning followed by extinction produces a memory that exhibits spontaneous recovery: behavior reverts back to the initially learned pattern. Spontaneous recovery is consistent with a mathematical model of learning in which memory is formed via multiple adaptive processes, one fast and the other slow. In this model, behavior shows spontaneous recovery because of the differing learning and decay properties of the two adaptive processes. Here, we identified motor commands that may reflect the contributions of these putative processes within a single saccade. Contrary to predictions of the current models, spontaneous recovery was not due to interacting dynamics of two putative processes. Rather, the commands during acceleration and deceleration periods adapted differentially, each with multiple timescales of learning.

In Experiment 1, subjects experienced a random horizontal error following conclusion of a vertical saccade. This error induced a change in the motor commands that guided the subsequent vertical saccade, partially correcting for the error. However, the error-dependent correction was not uniformly expressed throughout the saccade. Rather, the correction took place only via commands that arrived during the deceleration period. This raised the possibility that the putative fast process expressed its contributions during the deceleration period.

To test for this possibility, in Experiment 2 errors were always in the same direction, thus allowing for accumulation of trial-by-trial learning. The resulting saccades exhibited acceleration period commands that learned relatively little from error, and also exhibited little or no forgetting during the set breaks. In contrast, the deceleration period commands adapted with high sensitivity to error, and also suffered from a large amount of forgetting during the set breaks. While the perturbation block set breaks produced forgetting, the same set breaks during the error clamp blocks induced little or no forgetting. In addition, the acceleration and deceleration period commands decayed at similar rates during the error clamp blocks. These results illustrated that while in some respects the acceleration and deceleration period commands exhibited features of the putative slow and fast adaptive processes, they also had features that were inconsistent with this framework.

To resolve these discrepancies and better understand the mechanisms of spontaneous recovery, in Experiment 3 we presented an adaptation block followed by extinction. In the subsequent error-clamp block the motor commands during the acceleration and deceleration periods both showed spontaneous recovery. This was inconsistent with the predictions of the 2-state model.

To account for the richness of behaviors, including differing rates of learning from error, differing rates of forgetting due to set break, and similar patterns of spontaneous recovery in error-clamp trials, we considered a new model of learning. In this model, the acceleration and deceleration periods of a single saccade were adaptively controlled via independent learning processes. Spontaneous recovery occurred because each process was supported by the two timescales of learning, one fast and the other slow. To test our conjecture, we fit the model to the measured data in Exp. 2, and then used it to predict the data in Exp. 3. The results suggested that distinct controllers contributed to adaptation of acceleration and deceleration period commands, and each controller was supported by multiple timescales of memory.

### A new model

Our data suggested that acceleration and deceleration period commands of a saccade were controlled by distinct adaptive processes. This conjecture is consistent with recording and lesion data from the cerebellum. Neurons in the caudal fastigial nucleus of the cerebellum, which are critical for control of saccade trajectory, contribute differentially to acceleration and deceleration periods (Fuchs et al., 1993, 2010; Kleine et al., 2003; Ohtsuka & Noda, 1991) (but see (Sun et al., 2016)). This is particularly evident in experiments that have unilaterally inactivated a cerebellar deep nucleus (Buzunov et al., 2013; Goffart et al., 2003; Kojima et al., 2014; Robinson et al., 1993). For example, the right fastigial nucleus appears to specialize in adjusting the deceleration phase of rightward saccades, whereas during the same saccade, the left fastigial’s contribution is more limited to the acceleration period. Thus, the acceleration and deceleration periods of a saccade may be served by separate adaptive controllers in the cerebellum.

Fig. 8 presents a model that in principle reproduces the behaviors that we have observed. In this model, cerebellar Purkinje cells (P-cells) are organized based on their preference for visual error (Herzfeld et al., 2015). Their preference for error is expressed via turning of complex spike probability with respect to the direction of the visual error. When a saccade concludes and there is a rightward visual error (red arrow, Fig. 8), the result is an increase in the probability of complex spikes for P-cells on the left side of the vermis. These P-cells project to the left fastigial nucleus, and contribute to deceleration period motor commands. The same rightward error is the anti-preferred vector for P-cells on the right side of the vermis. These P-cells project to the right fastigial nucleus and contribute to acceleration period motor commands.

**Figure 8.**
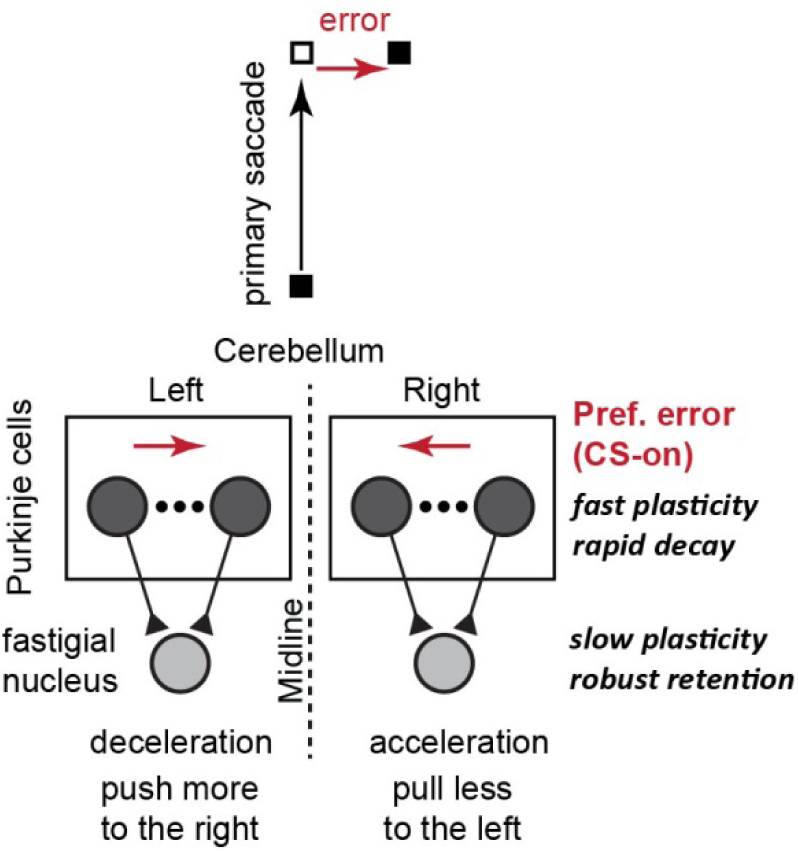
Working model of the cerebellum and its role in spontaneous recovery of saccades. Following production of a vertical saccade, there is a visual error: the target is to the right of the fovea. This produces unexpected activity in the left superior colliculus (and other regions), resulting in engagement of the contralateral inferior olive. The result is increased probability of complex spikes among the Purkinje cells (P-cells) to the left of the oculomotor vermis of the cerebellum. The same visual error results in suppression of the complex spikes among the P-cells to the right of the cerebellum. Both the presence and absence of complex spikes result in plasticity (LTD and LTP, respectively). This plasticity represents the fast time scales of learning. Fastigial nucleus cells receive inputs from the P-cells, and differentially contribute to the motor commands during the acceleration and deceleration periods. The inputs from the P-cells to the nucleus cell may also undergo plasticity. This plasticity represents the slow timescale of learning. Thus, distinct circuits contribute to acceleration and deceleration period commands of a single saccade, and each circuit learns from error via multiple timescales.

The rightward error increases the probability of complex spikes for the P-cells on the left of the vermis, but decreases this probability for the P-cells on the right. Both the presence and lack of a complex spike leads to plasticity (Herzfeld et al., 2018; Yang & Lisberger, 2014): depression of parallel fiber synapses follows presence of the complex spike, and potentiation follows its absence. On the next vertical saccade, there are reduced simple spikes that converge on the left fastigial, but increased simple spikes that converge on the right fastigial. Both result in increased force production to the right, but for the left fastigial nucleus, the expression of the forces is during the deceleration period, whereas for the right fastigial, the expression is during the acceleration period.

However, P-cell plasticity is not sufficient to account for the rich behaviors that we have recorded. Rather, learning of the acceleration and deceleration period commands must each be supported with a fast and a slow process (Fig. 7B). Thus, we speculate that there is plasticity elsewhere in this circuitry, possibly at the nucleus (Ohyama & Mauk, 2001). In the model, the P-cells exhibit plasticity that has high sensitivity to error, whereas nucleus cells exhibit plasticity that has low sensitivity. The two timescales of adaption for each period of the movement arise from distinct rates of plasticity in the cerebellar cortex (P-cells), and the nucleus.

The model provides a number of predictions. Following experience of an error, there is plasticity in both the P-cells that prefer that error (CS-on), and the P-cells for which this error is in the anti-preferred direction (CS-off). There is some evidence for this conjecture. In saccade and pursuit tasks it has been shown that presence of a complex spike in one trial is followed by adaptive changes in simple spikes that P-cells produce in the subsequent trial (Herzfeld et al., 2018; Medina & Lisberger, 2008; Yang & Lisberger, 2014). Similarly, lack of a complex spike is also associated with changes in the number of simple spikes and trial-to-trial changes in behavior (Herzfeld et al., 2018; Yang & Lisberger, 2014). Our model predicts that acceleration period commands learn slower because lack of complex spikes will cause less learning than presence of complex spikes. Prior work has found evidence for this differential rate of plasticity (Herzfeld et al., 2018; Yang & Lisberger, 2014), but the prediction remains to be tested.

The model predicts that spontaneous recovery is because of differential rates of learning in the cerebellar cortex and nucleus. This is a more speculative component of our model. However, in a classical conditioning task there is evidence that behavior improves because of plasticity in both the cerebellar cortex neurons and the nucleus neurons, but the plasticity may be slower in the nucleus neurons (Ohyama & Mauk, 2001).

Finally, the model predicts that time-dependent decay of plasticity should be larger among the neurons in which learning is faster, i.e., P-cell plasticity should decay faster than plasticity induced in the nucleus cell. These predictions can be tested by recording from the two regions during trial-by-trial adaptation.

Does this model generalize to other kinds of movements? The idea that there may be separate adaptive controllers for acceleration and deceleration may be relevant to reaching. During reaching, many neurons in the interposed nucleus of the cerebellum tend to be active during the deceleration period of arm extension, whereas other neurons are active during the acceleration period (Becker & Person, 2019). Stimulation of the nucleus neurons during arm extension produces a premature termination of the reach, resulting in over activation of flexion muscles during the deceleration period. Thus, distinct cerebellar deep nucleus neurons may contribute to acceleration and deceleration commands during reaching.

However, single trial learning in reaching exhibits an error-dependent response that is different than the one we found during saccades: in saccades, learning from a single error results in corrective motor commands that are expressed primarily during the deceleration period (Fig. 1). In reaching, experience of an error produces corrective motor commands on the subsequent trial that are expressed throughout the reach (Albert & Shadmehr, 2016). Thus, while the cerebellum may contribute differentially to acceleration and deceleration periods of reaching (Becker & Person, 2019), we have not observed behavioral evidence for this dissociation during reaching (Albert and Shadmehr, 2016).

Overall, our results suggest distinct adaptive controllers contribute to the acceleration and deceleration phases of a single saccade, and that each controller is supported by a fast and a slow learning process.

## Acknowledgements

The work was supported by grants from the National Science Foundation (CNS- 1714623), the NIH (R01-NS078311, R01-NS096083), and the Office of Naval Research (N00014-15-1- 2312). SPO was supported by a fellowship from the NIH (1F31-NS108731).

